# Hepatic fibrosis induced zinc-deficient dermatosis in American alligators (*Alligator mississippiensis*)

**DOI:** 10.1101/2022.08.30.505809

**Authors:** Ilaria M. Piras, Annemarie Bezuidenhout, Josué Díaz-Delgado, Deirdre Slawski, Pamela A. Kelly

## Abstract

Crocodilian farming generates strong economic incentives for the conservation of several species previously endangered by intensive hunting. Ranching farms, in particular, are intimately connected to the natural crocodilian habitat and have a significant impact on wetland preservation. The financial sustainability of this industry relies on the production of first grade skins for the luxury leather market. Only flawless skins are considered of first grade by the stringent standards of the market, and even a single defect represents an economical loss. “Double scale” is one such defect that drastically reduces the appeal of crocodilian skin. Although double scale defects represent a threat to the economical sustainability of the farming industry, there is no scientific literature available on this topic. This study, carried out in a ranching farm of American alligators (*Alligator mississippiensis*), represents the first investigation into the pathogenesis of double scale. Our results indicate that double scale is a keratinization disorder associated with zinc deficiency. Furthermore, we found that portal hypertension due to liver fibrosis, underlies zinc deficiency in cases of double scale. Lastly, we found that chronic vitamin A toxicity can cause liver fibrosis in crocodilians. For the first time, we demonstrate a causal association between liver disease and skin quality in a crocodilian species. This study reveals the conserved role of zinc in the homeostasis of reptilian skin. Also, we show that, like mammals, reptiles may develop liver fibrosis following chronic vitamin A toxicity and through activation of hepatic stellate cells. Our results advance herpetological medicine and will translate into improved captive crocodilian welfare and husbandry.

## Introduction

After decades of unregulated and intensive hunting for skin exploitation, most crocodilian species became endangered, and some were almost extirpated by the beginning of 1970s^1^. This dramatic decline of wild crocodilian populations induced several countries to protect crocodilians through legislation. In addition, the worldwide trade of wild species and their products became regulated by the Convention on International Trade in Endangered Species (CITES) from 1975. Crocodile farming gained momentum in this historical context as a valuable opportunity to provide skins for an ever-growing leather market and, at the same time, incentivize the conservation of crocodilians^2,3^.

Ranching is the most common way of farming American alligators (*Alligator mississippiensis*), in the United States. Ranching farms are intimately connected to the natural alligator habitat and have a significant impact in wildlife conservation. This way of production is based on the collection of wild eggs, followed by regular reintroduction of juveniles and hatchlings born on farm, to maintain the wild crocodile population^4^. This practice incentivize the preservation of the wet land habitats, saves eggs that are often subject to predation or destruction in wild conditions^5^ and boosts hatchlings survive until adulthood^5,6,7,8^.

The main focus of crocodile farming is to produce high quality skins to supply the expanding demand for premium hides for the luxury leather market^7^. For this reason, care is taken during rearing to minimize damage to the belly skin, which is the most valuable part of the hide, either from abrasive surfaces, from interactions with other crocodiles, and from any dermatologic conditions that can produce a defect^4,9,10^.

This study is a clinical and pathological investigation conducted on a ranching farm of American alligators following a year of poor skin gradings due to high incidence of “double scale” (DS) defects in the population. This skin condition reduces leather quality and has an important impact on the financial sustainability of the industry. The manifestations of this condition remain uncharacterized, and the etiology is unclear. Multifactorial causes, involving potential nutritional, metabolic and genetic factors have been considered anecdotally, but none have been explored in the scientific literature so far.

## Results and discussion

### Double scale is a disorder of keratinization

The skin is a protective barrier composed of multiple layers of specialized epithelial cells called keratinocytes, that offer protection against pathogen invasion, chemical, thermal, and physical damage, and prevent body water loss. All these functions are provided by the outermost layer of the epidermis composed of terminally differentiated keratinocytes, the corneocytes. The constant balance of keratinocyte proliferation, differentiation and shedding of corneocytes facilitates repair after external trauma and maintains the steady state of the corneal layer^11^. Scales are skin appendages evolved by reptiles^12^, which exquisite arrangement pattern has made crocodilian leather one of the most demanded products of the luxury fashion market since 1800^1,13^.

DS defect undermines the appeal of the belly scale pattern with an array of defects. In this study, DS presented as a combination of focal extensive pitting and roughening of the cranial and medial edge of the belly scale (18/18 -100%), which was often accompanied by a brown discoloration (Figure 1A). In the most severe cases, a linear, ring-shaped dent carving the central portion of the scale was also present (7/18-38.8%) (Fig. 1A, arrow). The presence of this dent is responsible for the duplicated appearance of the scale, from which this defect derives its name.

**Figure 1.**
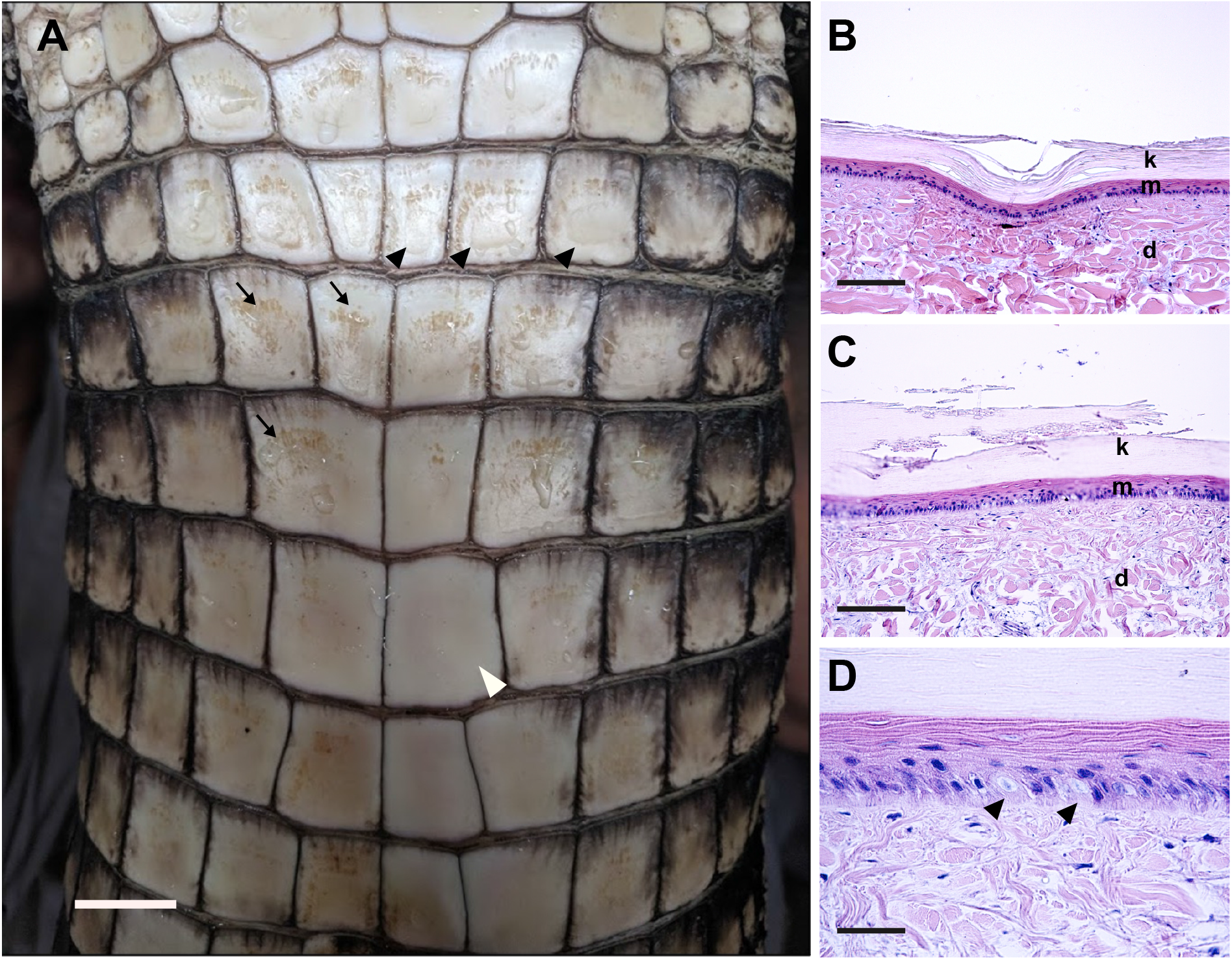
Belly skin with double scale defect, American alligator. Gross appearance of double scale defect (A) characterized by a rough and pitted surface (arrows) often associated with a linear dent (black arrowheads). Note the smooth and even surface of the normal scale (white arrowhead). Scale bar 1 cm. Microphotographs of a cross section of double scale (B-C-D). H&E, scale bar 200um (B-C), 50um (D). The grossly noticeable dent is histologically characterized by a focal depression of the epidermis and the dermis (B). At the gross level of the rough surface, histologically the corneal layer is disrupted and fragmented (C). Higher magnification of the epidermis shows frequent degeneration of the basal keratinocytes (arrowheads) (D). (k) corneal layer and (m) Malpighian layer of the epidermis, (d) dermis.

Histologically, this dent presented as a depression of the epidermis and superficial dermis (Figure 1B) giving the scale an irregular profile that resulted in downgrading of the processed skin (data not shown). DS also presented loss of the characteristic compactness of the reptilian β-keratinized corneal layer, with frequent multifocal fragmentation (Figure 1C) that corresponded grossly to the roughened pitted areas of the scale. Other changes evident at histological examination were increased numbers of degenerating basal keratinocytes with a mean of 3.7 per 10,000 cells, (SD=9.3, n19) (Figure 1D) compared to 0.21 per 10,000 cells in normal scales (SD=0.39, n=9). β-keratin layer thickness was 94.6 µm on average in normal scales (SD=11.6, n= 18) (figure 2A, 2B and 2G), which is consistent with measurements reported in the literature for juvenile alligators of similar age^14^. The α-keratin, measured at the level of the hinge, was 18.8 µm on average (SD=5.3, n=18) (Figure 2C and H). A marked thickening of the keratin layer without retained nuclei, or orthokeratotic hyperkeratosis (Figure 2), characterized all DS samples examined (18/18-100%) compared to normal scales (n=9) (Figure 2A and 2B). This hyperkeratosis affected both, the β-keratin layering of the scale plate (corneal layer, mean= 168.5µm; SD=35.4, n=36, *p* value <0.0001) (Figure 2D and G), and the α-keratin in the hinge area (mean= 70.5 µm; SD=21.5, n= 36 *p* value <0.0001) (Figure 2E and 2H). The thickness of the live strata of the epidermis, including the basal, supra-basal and transitional layers (also referred to as Malpighian layer), was significantly reduced in scales with hyperkeratosis (average of 33.1 µm, SD=9.9 µm, n=36, *p* value <0.0001) compared to normal scales (mean 51, SD=11.9, n18) (Figure 2B, 2D and 2F). In addition, changes to the dermis were evident. Underlying the epidermis, the dermis is the skin stroma composed of collagen bundles, ground substance, blood vessels and nerves that provides innervation, vascularization, and support to the organ.

**Figure 2.**
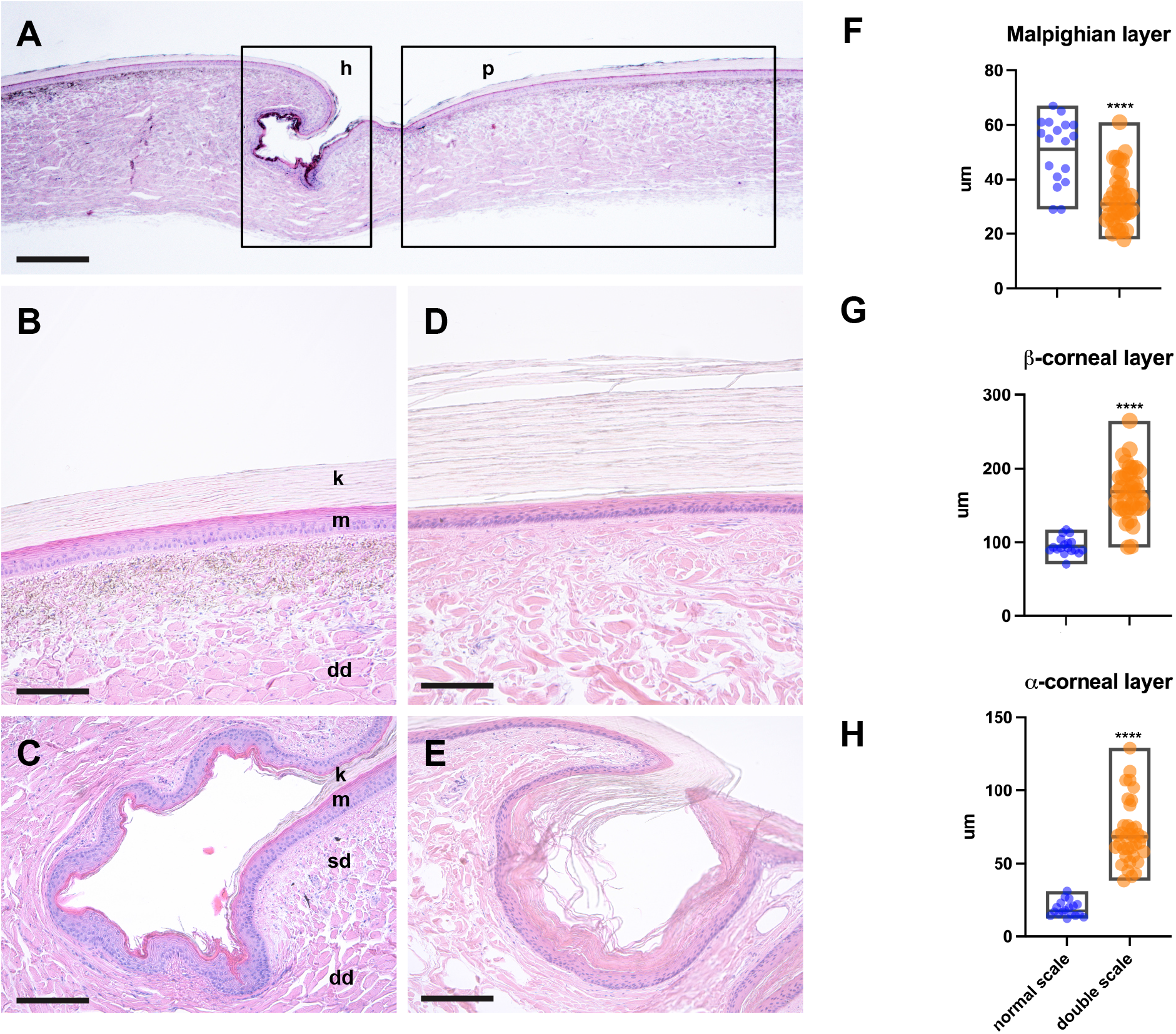
Hyperkeratosis characterizes double scale in American alligator. Microphotographs of a normal scale (A-B-C) and double scale (C-D). H&E. Normal anatomy of alligator scales (A) showing the plate region (p box) constituting most part of the scale and the hinge region (h box) which connects the cranial (left) to the caudal (right) scale. Scale bar 1mm. Compared to normal scale (B and C), the keratin layer of skin affected with double scale is markedly thickened both at the level of the plate (D) and the hinge (E). Note the β-keratin covering the plate of the scale stains negatively (B and D) whilst the α-keratin at the level of the hinge dyes pink with eosin (C and E). The live strata of the epidermis, also called Malpighian layer, is instead thinner (E). Scale bar 200µm. (k) Corneal layer, (m) Malpighian layer of the epidermis, (sd) superficial dermis, and (dd) deep dermis. The box graphs show measurements of Malpighian layer (F) beta (G) and alpha (H) corneal layers of normal (n=18 measurements) and double scales (n=38 measurements). Central line in the box indicates mean. Difference of means calculated with Student *t* test was significant with *p* value <0.0001 (****).

A disarray in the orientation of collagen fibers was evident in the superficial and deep dermis of several DS individuals (9/19; 47%), accompanied by a multifocal increase of the ground substance (Figure 3). Disarray and loss of collagen fibers could be considered responsible for the altered staining properties noted in processed alligator skin with DS defect (data not shown).

**Figure 3.**
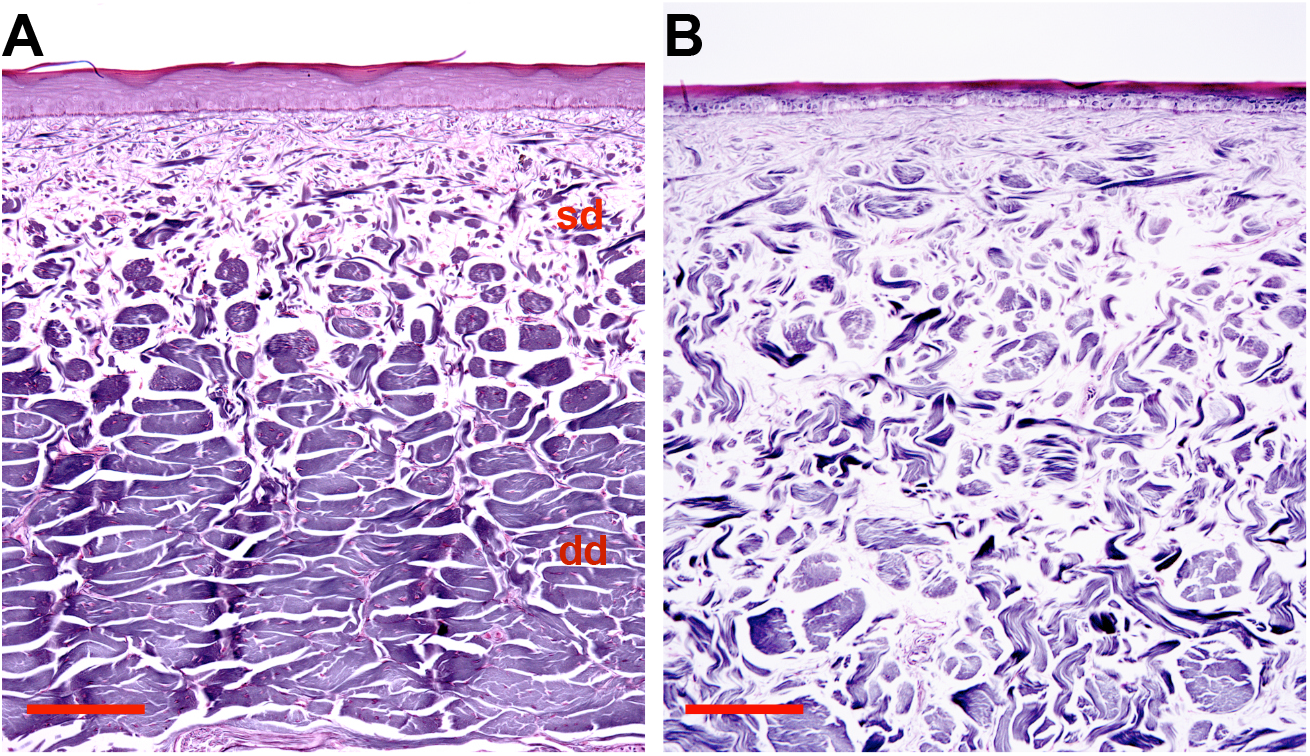
Dermal collagen disarray characterizes the skin with double scale. Microphotographs of a normal scale (A) and double scale (B). Reticulin Stain. Scale bar 200µm. Normal anatomy of alligator dermis (A) showing and orderly arrangement of collagen bundles (stained black) that became larger in the deep dermis. Note that all bundles are oriented perpendicular to the cut section the separation between superficial and deep dermis is well-defined. In double scales (B) collagen bundles are reduced in size and are separated by an increase of clear spaces (ground matter). The bundles are arranged haphazardly and a clear definition between superficial and deep dermis is lost.

### There is a correlation between skin lesions and liver injury

A further, systematic histopathological examination of intestine, lung, heart, brain, spleen, liver, and kidneys from 86 alligators revealed a high incidence of liver disease (47/86; 54%); 70% of pathologic livers (33/47) presented fibrosis with variable severity (Figure 4A and Supplementary Figure 1). Most interestingly, liver fibrosis was always present in alligators with DS (18/18; 100%). No other related histopathological findings were observed in other tissues. There is still little information available on normal liver histology of reptiles, and to the best of our knowledge, liver fibrosis has not been described in alligators. Hepatic fibrosis represents a scarring response to chronic liver injury after a variety of insults^15^. The most advanced stage of fibrosis, called cirrhosis, defines the distortion of the liver parenchyma with bridging scars and nodule formation, accompanied by altered blood flow^15^. In the alligator livers examined, fibrosis formed within and around periportal tracts and the liver capsule. In the most severe cases, fibrosis bridged several portal areas resulting in complete loss of normal lobular architecture with evidence of nodular regeneration, as described in cirrhosis of mammalian species (Figure 4C). Other liver changes consistently associated to fibrosis in alligators were bile duct proliferation (Figure 4A), which is considered a nonspecific reaction to liver injury described in humans and several domestic species,^16^ glycogen accumulation (figure 4E), causing hepatocytes vacuolation and swelling and increase of periportal melanomacrophage numbers (Figure 4A). Except for the expected accumulation prior to hibernation, glycogen accumulation is largely considered a consequence of hepatocytes metabolic pathways disturbance^17^. An increase of melanin laden macrophages (melanomacrophages) suggests an enhanced phagocytic activity in reptilians^18^ and is likely due to an increased rate of cell debris clearance, resulting from hepatocyte loss in these cases. Myofibroblasts are considered the main source of collagen production in liver fibrosis in humans, mice, and dogs^19,20^. They are fibroblast-like cells that express α-smooth muscle actin which confers them contractile properties^19,17^. As expected, immunolabelling of alligator livers showed α-SMA-positive cell numbers were markedly increased around periportal areas, where the collagen accumulated (Figure 5A).

**Figure 4.**
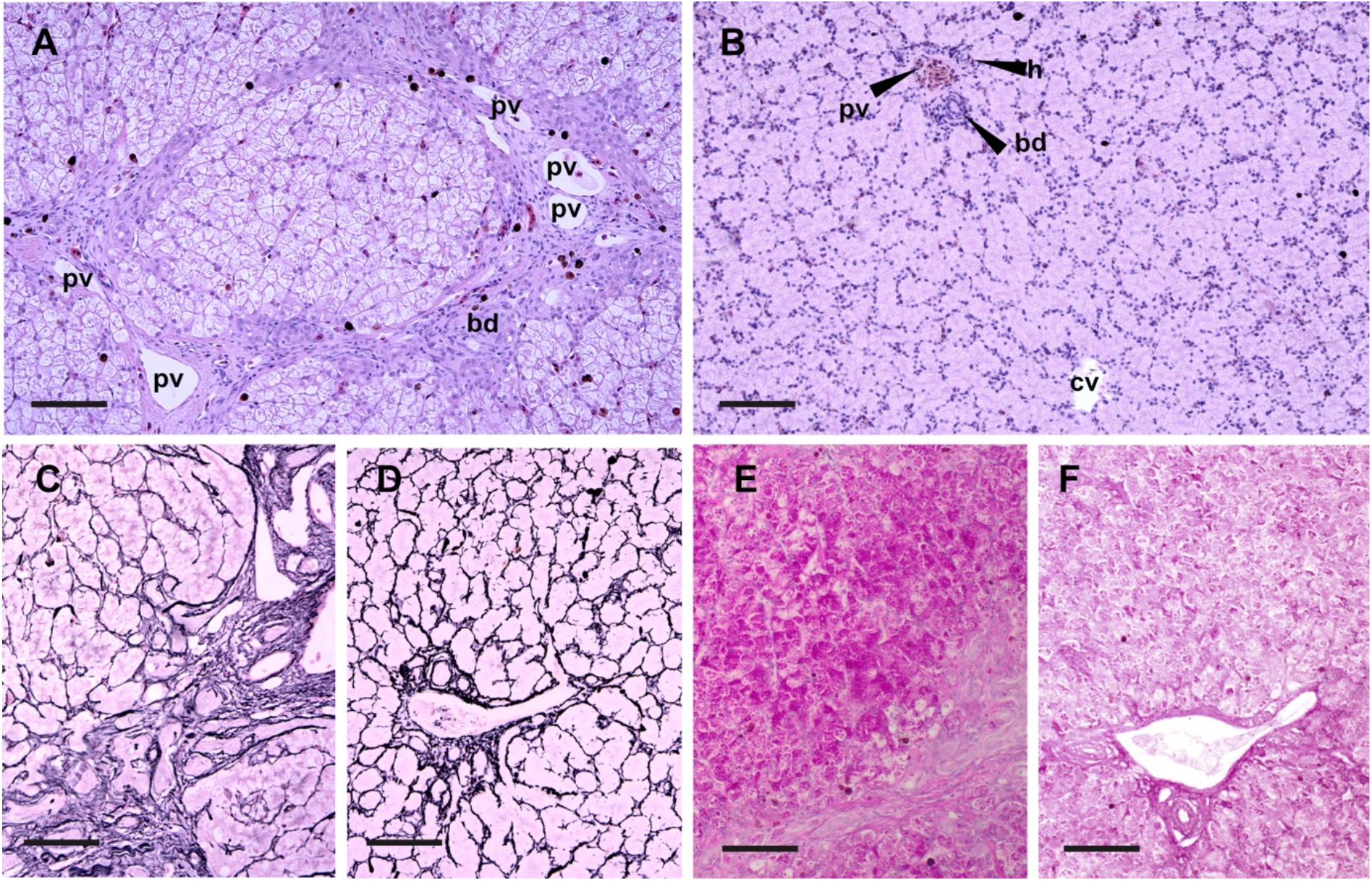
Liver fibrosis in American alligators with double scale. Microphotographs of alligator livers (A-B) H&E. (C-D) Reticulin stain. (E-F) PAS. Scale bar 200µm. Liver affected by severe fibrosis (A) showing bands of collagen fibers bridging between several triads associated with increased numbers of melanin laden macrophages. Biliary ducts are increased in number and are tortuous. Portal veins were tortuous and irregularly dilated, suggestive of shunting. Hepatocytes show diffuse feathery vacuolation and swelling. Central vein is collapsed and undetectable. Normal liver lobule architecture (B) showing a triad (upper left) composed of one portal vein, one hepatic artery and one biliary duct. A central vein (lower right) is seen at the center of the lobule. Reticulin stain accentuates the nodular deformation of the lobule and distortion of hepatocellular plates and nodular deformation of hepatic parenchyma (c) in cirrhotic liver compared to a normal one (D). Periodic acid Schiff stain highlights glycogen accumulation is causing hepatocyte vacuolation of livers with fibrosis (E) compared with normal livers (F). Biliary ducts (bd), portal vein (pv) and (h) hepatic artery and (cv) central vein.

**Figure 5.**
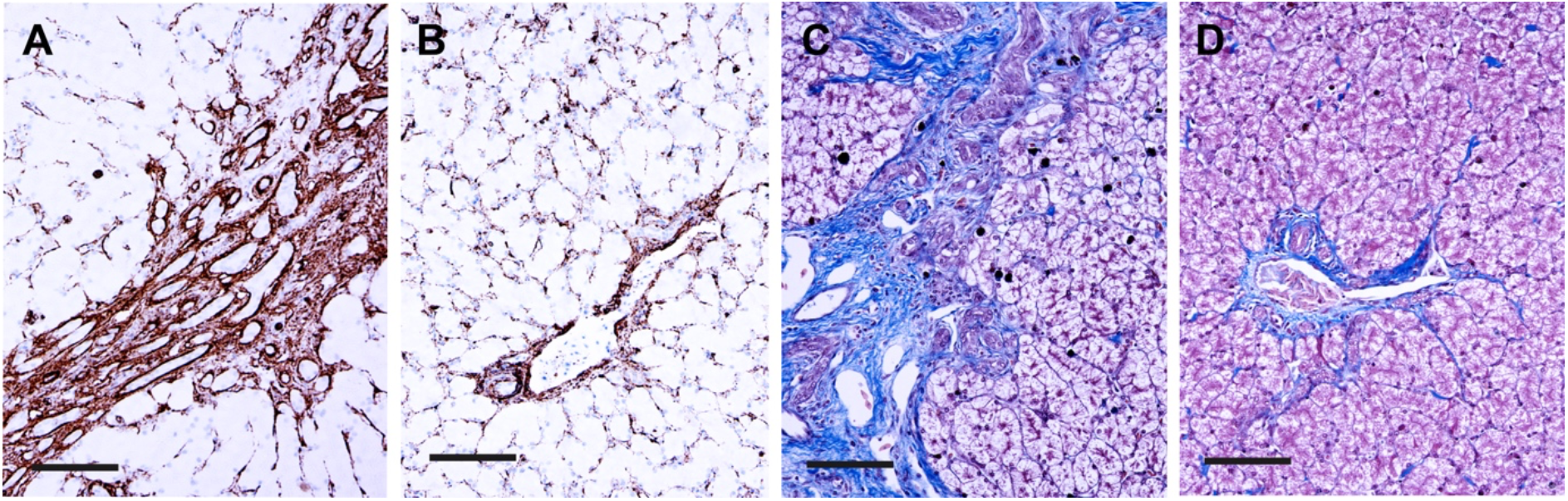
Myofibroblasts are the fibrogenic cells in American alligator’s livers. Microphotographs of alligator livers. (A-B) Anti-α-smooth muscle (SMA) immunohistochemistry. (C-D) Masson’s Trichrome. Scale bar 200µm. In livers with fibrosis α-SMA-positive myofibroblasts increase within periportal areas (A) in association with collagen deposition (blue stain, C). Note the numbers of myofibroblasts in the perisinusoidal spaces are reduced compared to normal livers (B). Collagen distribution in normal livers (D).

Myofibroblasts in livers can be recruited by activation of hepatic stellate cells (or Ito cells), liver resident fibroblasts (portal or centrilobular), epithelial cells that undergo epithelial-to-mesenchymal transition, bone marrow-derived fibrocytes, and smooth muscle cells that surround blood vessels^21^. Comparison of α-SMA and desmin immunolabelling with normal alligator livers indicated hepatic fibrogenic cells derived from proliferation of periportal and perivascular fibroblasts but also recruitment of desmin positive cells from the sinusoids, which are most likely hepatic stellate cells (Figure 5 and 6).

**Figure 6.**
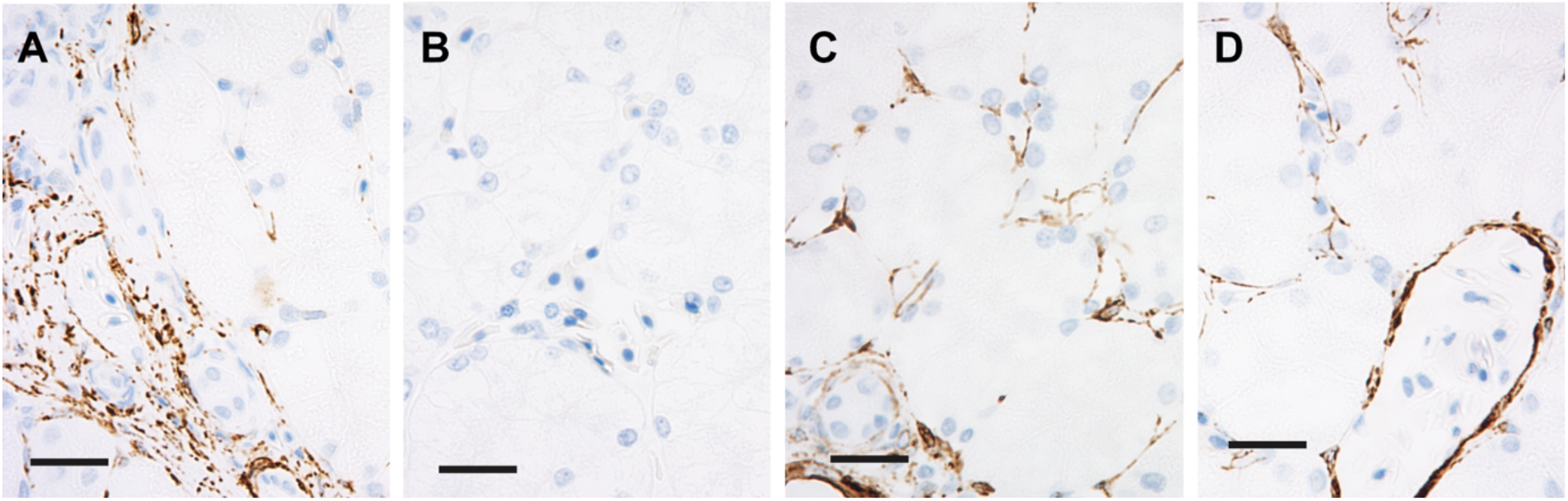
Migration of desmin-positive cells from perisinusoidal spaces to periportal areas in association to liver fibrosis in American alligator. Microphotographs of alligator livers. Anti-desmin immunohistochemistry. Scale bars 50µm. When fibrosis is present immunolabelled cells are concentrated in the periportal areas (A), leaving the sinusoids depleted of desmin-positive cells in the midzonal and centrilobular areas (B). In normal livers, desmin-positive cells have cytoplasmatic processes that diffusely and evenly stretch into the perisinusoidal spaces in both periportal (C) and centrilobular areas (D).

### Vitamin A toxicity underlies liver injury

Progressive liver fibrosis can be caused by chronic viral infection, prolonged use of hepatotoxic drugs, longstanding exposure to aflatoxin, iron, and copper accumulation, chronic biliary cholangitis, autoimmune hepatitis, and chronic vitamin A toxicity among others^19,22,23^. Alcohol abuse, non-alcoholic fatty liver disease, non-alcoholic steatohepatitis are largely reported in humans^16^.

Results from hepatic vitamin and mineral level analysis (table 1) indicated the average concentration of vitamin A in alligator livers with fibrosis was 880,000 mg/100g (n= 27). This value is significantly higher than vitamin A concentration found in normal alligator livers (389,000 mg/100g; n=22) examined in this study. In our alligators, hypervitaminosis A resulted from a longstanding diet with vitamin A concentration ranging from 23,300 IU/Kg to 38,200 IU/Kg. The tolerable range of dietary vitamin A levels for crocodilians is currently unknown. Vitamin mix supplementation is generally used in excess in commercial crocodilian diets as no toxicities have been reported in the literature, so far^24^.The maximum tolerable dose of dietary vitamin A for chicken broiler breeders appears to be 35,000 IU/kg, corresponding to 195.2 mg/100 g of retinyl esters in the liver whilst higher supplementation has been shown to impair liver function^25^. Compared to chickens and turkeys, which are the closest phylogenetically related domestic species, it seems that alligators have a much higher capacity to accumulate vitamin A in the liver^25,26^. Considering that 70% of alligators fed with vitamin A concentration above 23,300 IU/Kg developed liver fibrosis, we suggest these levels may be toxic. Liver damage related to chronic hypervitaminosis A is a rare, but well described condition in humans^27^ and has only been reported once in a cat^28^. Although occasionally suspected in pet reptiles, it has not been characterized^29^. The pathologic changes in the liver are related to the increased retinyl storage in the organ and may result in portal hypertension, secondary to sinusoidal compression, and hyperplasia of Ito cells followed by fibrosis^16,30^. Toxicological studies are warranted to confirm the toxic ranges of vitamin A in crocodilians.

**Table 1.** Micronutrient concentration in liver and serum of alligators with liver fibrosis compared to alligators with normal livers. Micronutrients in the liver and serum are indicated as mean value in µg/g and µg/mL respectively. Means are calculated on 22 samples from animals with normal livers and 28 with liver fibrosis. Difference of means was calculated with Student *t* test. Significant results with *p* value <0.0001 are marked as ****, *p* value 0.0021 as **, and *p* value <0.05 as*.

|  | Controls |  | Liver Fibrosis |  |  |  |  |  |
| --- | --- | --- | --- | --- | --- | --- | --- | --- |
|  | Liver | Serum | Liver | p value | Trend | Serum | p value | Trend |
| **vitamin A** | 389 |  | 880 | **** | ↑ |  |  |  |
| **Palmitate** | 362 |  | 855 | **** | ↑ |  |  |  |
| **retinol** | 28 |  | 41 |  |  |  |  |  |
| **Iron** | 388 | 136 | 734 | **** | ↑ | 182 |  |  |
| **Zinc** | 81 | 1 | 96 | * | ↑ | 0.6 | **** | ↓ |
| **Copper** | 39 | 0.67 | 57 |  |  | 0.55 | * | ↓ |
| **Selenium** | 2.9 | 324 | 3.3 | * | ↑ | 353 |  |  |
| **Manganese** | 5.4 | 8 | 3.4 | **** | ↓ | 10 |  |  |
| **Molybdenum** | 0.29 | 19 | 0.13 | **** | ↓ | 22 |  |  |
| **Cobalt** | 0.02 | 0.2 | 0.02 |  |  | 0.7 |  |  |
| **Arsenic** | 0.09 | 0.017 | 0.1 |  |  | 0.025 | ** | ↑ |
| **Cadmium** | 0.02 | 0.1 | 0.02 |  |  | 0.1 |  |  |
| **Thallium** | 0.02 | 0.1 | 0.02 |  |  | 0.1 |  |  |
| **Lead** | 0.046 | 0.1 | 0.023 |  |  | 0.1 |  |  |
| **Vitamin E** | 69 | 4.9 | 57 |  |  | 3.4 | ** | ↓ |

Hepatic iron levels were also increased in alligator livers with fibrosis (Table 1). Histologically, iron increase and pattern of accumulation occurred exclusively within periportal macrophages (Supplementary figure 2). The mesenchymal pattern of iron accumulation together with the lack of stainable iron in hepatocytes and other tissues indicates the origin of iron loading is secondary to hepatocyte loss rather than chronic, augmented intake^31^. Other causes of liver disease leading to fibrosis such as copper loading, aflatoxin and CCl_4_ exposure were excluded based on the result of liver oligoelements analysis and toxicologic test of feed and water. As histological evidence of cholangiohepatitis and bacterial infection were absent in the livers examined, they were discarded as possible underlying causes of liver disease.

### Zinc deficiency due to portal hypertension is the cause of skin lesions

Mineral and vitamin analysis of the serum and livers revealed an imbalance between hepatic accumulation and systemic distribution of several elements in alligators with liver fibrosis (Table 1). Specifically, vitamin E, copper and zinc were significantly decreased in peripheral blood, although their hepatic concentration remained steady in the liver or, like zinc, slightly increased (Table 1). These imbalances suggested portal hypertension had developed in these individuals, which is supported by histopathological observations of sinusoidal compression, myofibroblast hyperplasia, sinusoidal and periportal fibrosis. All these changes in alligator livers, most likely contributed to increased intrahepatic blood flow resistance, impairing the systemic availability of nutrients absorbed by the gut and delivered to the liver through the portal vein.

The venous shunting seen in histological sections as tortuous and irregularly dilated portal veins in livers with fibrosis (Figure 4A, asterisk) is an attempt to correct the circulatory disturbances present in the organ and is consistently present in human livers with portal hypertension^32^.

Of all serum micronutrient imbalances in alligators, zinc reduction was the most striking for its severity (0.6 ug/mL; n= 20) compared to controls (mean 1µg/mL; n= 22) (Figure 7D). These results are highly suggestive of zinc deficiency as a primary cause of alligator skin lesions, specifically DS. Zinc deficiency is a common finding in people suffering of liver cirrhosis and is due to portal hypertension or decreased serum levels of albumin^33^. In this case albumin serum levels of alligators with liver disease were comparable to the controls (1.77 and 1.8 g/dL, respectively). Cutaneous lesions are common manifestations of zinc defficiency both in humans and in animals, but the pathogenic mechanisms are unclear^34^. Zinc in the skin is regulated by transporters (ZnTs and ZIPs) and metallothioneins (MTs) which regulate epidermal proliferation and differentiation. MTs are cysteine-rich, zinc-binding proteins synthetized in response to tissue zinc levels and essentially control its availability^35^. MTs also act as free radical scavengers^36^. Several authors suggest that zinc antioxidant effects explain its inhibitory effect on apoptotic pathways, which is considered a key aspect of the pathology of zinc-defficient skin^37^.

**Figure 7.**
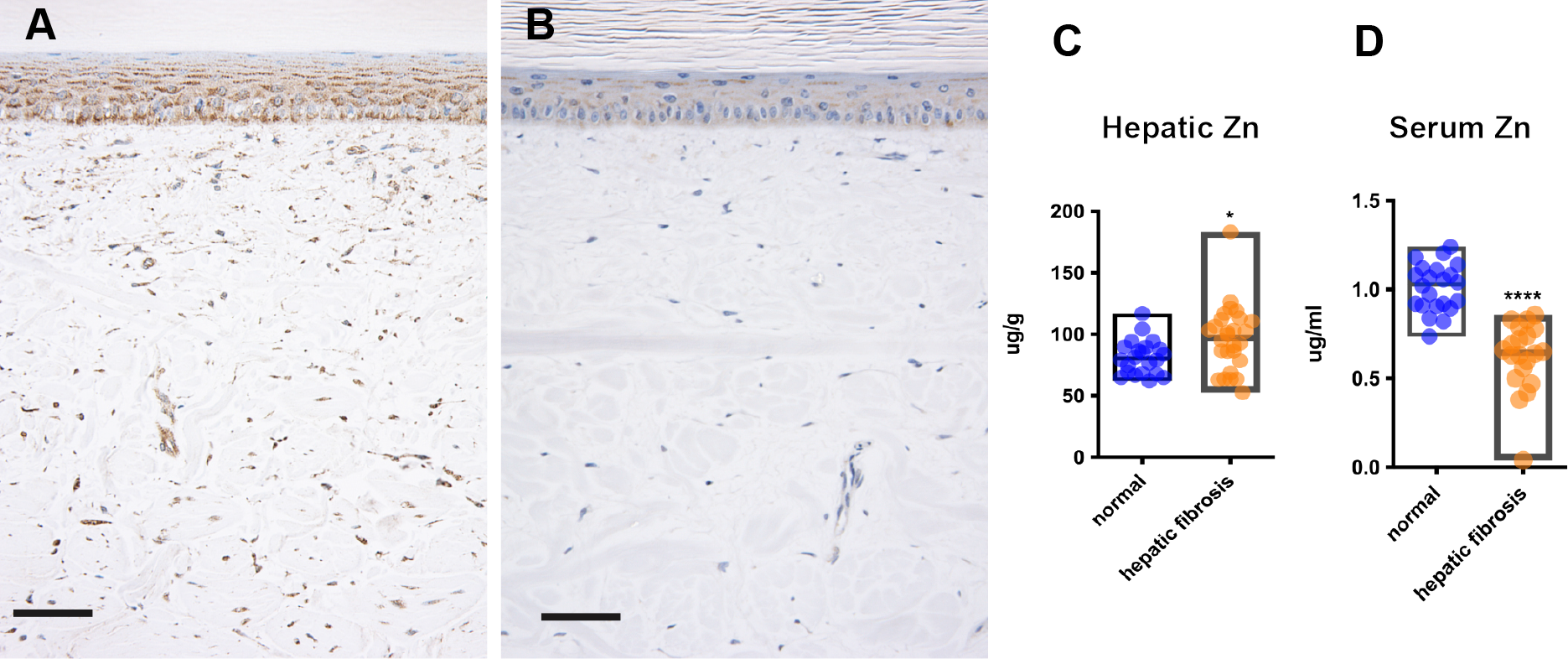
Zinc deficiency and metallothionein depletion in the epidermis of alligators with liver fibrosis. (A-B) Microphotographs of belly scale. Anti-metallothionein immunohistochemistry, scale bar 50µm. Metallothionein is labelled (granular cytoplasmic pattern) diffusely in all layers of the live epidermis and dermal fibroblasts in normal scales (A). Very scant metallothionein immunolabelling is detectable in the epidermis and dermis of animals with double scale (B). Box graphs show levels of hepatic (C) and serum (D) zinc in alligators with liver fibrosis (n=28) compared to alligators with normal livers (n=22). Central line in the box indicates mean. Difference of means calculated with Student *t* test. Significant differences are indicated with **** for with *p* value <0.0001 and * for *p* value <0.05.

MTs expression may be used for subjective determination of zinc concentration in skin samples^34^. Anti-metallothionein immunolabelling of alligator skin revealed a drastic reduction of MTs in alligators with liver fibrosis, confirming zinc deficiency is a primary underlying cause of skin lesions in these animals (Figure 7).

The presence of increased numbers of degenerated basal keratinocytes in the skin samples examined seem to support the presence of an increased oxidative stress in the epidermis of alligators. Interestingly, the histopathology of zinc-deficient dermatosis in alligators seems to differ from humans and domestic animals. In mammalian species the characteristic thickening of the corneal layer is accompanied by the retention of nuclei in corneocytes (parakeratosis) and the layers of live strata of the epidermis are also increased (hyperplasia or acanthosis)^34^. These morphological differences are likely due to the different mechanism of keratinization in crocodilian skin, where terminally differentiated keratinocytes produce β-keratin, which is exclusive of reptilian and avian appendages and is responsible for the rigidity of scales and feathers. Another important difference concerns keratohyalin granules, which are essential in mammalian keratinization process and absent in crocodilian keratinocytes. Consistent to all species with zinc-deficient dermatosis, is the severe thickening of the corneal layer which is considered secondary to disorders of proliferation, differentiation, and exfoliation of the epidermis^34^, although the precise pathogenesis remains unknown. Regarding the collagen disarray present in the dermis of scales with DS defect, we suggest an altered function of zinc-dependent matrix metalloproteinases could be involved. Matrix metalloproteinases belong to a family of endopeptidases involved in the cleavage of extracellular matrix proteins, required for tissue remodeling and wound healing^38^. We hypothesize that a lack of tissue remodeling capability would cause the collagen disarray in alligator skin with zinc deficiency. Further research is warranted to assess this hypothesis.

## Conclusions

This study reveals how liver disease can affect skin quality in farmed alligators, highlighting the importance of animal health and welfare for the economical sustainability of crocodilian farming. Double scale is considered anecdotally as a multifactorial skin condition; here we demonstrate how zinc deficiency due to hepatic portal hypertension plays a main role. This finding indicates a conserved role of zinc in reptilian skin homeostasis, suggesting that acquired nutritional and metabolic disturbances can lead to skin disease in this species. There is limited knowledge on nutritional needs of captive crocodilians and literature lacks reference ranges. The liver disease in this case was likely due to chronic vitamin A toxicity. Based on histopathological examination of livers, we suggest dietary concentrations of vitamin A higher than 23,000 IU/kg are potentially hepatotoxic, whilst an average of 15,000 IU/kg seem to be safe for captive alligators.

## Methods

### Ethic statement

This study was ethic review exempt (AREC-E-22-20-KELLY) as it involves samples collected exclusively during normal husbandry or veterinary clinical procedures. No procedures were carried out on live animals and no animal was euthanized for the sole purpose of this study.

### Animals

All alligators included in this study were reared on the same farm. At the time of investigation, the farm produced over 12,000 animals per annum on a single site. Animals were housed in groups of 20 to 80 depending on their size. These groups consisted of a mix of both male and female alligators. The houses were insulated and were divided into pools and feeding and basking decks. Diet consisted of a commercially available complete crocodilian pelleted feed purchased from a single manufacturer that was tested routinely for mycotoxins.

### Sample Collection

Between April 2019 and June 2020, during routine abattoir procedures, samples of liver, lung, kidney, heart, spleen, intestine, brain, and serum were collected from 86 randomly selected alligators (Supplemental table 1). From the same animals we collected 18 samples of belly skin with double scale and 9 samples of normal belly skin (Supplemental table 1). Age range for selected animals was between 9 and 19 months; sex and farming pen of origin were mixed. The average length and belly width for each age group of alligator at harvest is presented in Supplemental table 2.

### Histopathology

Following fixation in 10% neutral buffered formalin, tissues were processed and then embedded in paraffin-wax, sectioned at 4 µm, and stained with hematoxylin and eosin (H&E). Histological sections were assessed for the presence of histopathological changes by pathologists P.A.K (ECVP board certified), J.D.D. (ACVP board certified) and I.M.P. (anatomic pathology resident).

3 representative samples of liver with fibrosis and 3 with normal histology were stained with Masson trichrome, Gordon and Sweet’s reticulin, Perls’ Prussian blue stains to highlight collagen, reticular fibers and iron respectively. To characterize the vacuolar hepatopathy further, Periodic acid Schiff (PAS) staining was also carried out on these samples.

The histopathological examination of skin was based on 61 scales with defects sampled from 18 alligators and 32 normal scales sampled from 9 individuals, for a total of 93 scales. Skin measurements were carried out on 18 normal and 36 scales with DS (2 scales per alligator) using imageJ on microphotographs of 20x magnification with scale set to 15 µm equivalent to alligator erythrocyte wide diameter.

Six representative samples of skin, three normal and three with DS defect, were also stained with Gordon and Sweet’s reticulin, Masson trichrome, PAS, Gram and used for immunohistochemical labelling.

### Immunohistochemistry

Fibrogenic cells were assessed by immunohistochemistry (IHC) on six samples of liver, three with fibrosis and three normal, using an anti-desmin antibody at concentration of 1/25 (a-desmin clone D33, M0760; Denmark A/S) and a monoclonal antibody directed against the alpha isoform of smooth muscle actin at a working dilution of 1/100 (a-SMA, clone 1A4, n M0851; Dako, Denmark A/S). Alligator skeletal muscle was used as positive control for anti-desmin antibody (Supplemental figure 3A). The tunica muscularis of alligator intestines served as positive control for a-SMA antibody (Supplemental figure 3C).

The presence of zinc deficiency was assessed targeting metallothionein using a rabbit polyclonal anti-metallothionein at 1/400 dilution (ab192385, abcam) on six samples of alligator skin, three with double scale defect and three normal. Mouse skin was used as positive control for the primary antibody (Supplemental figure 3E). For all antibodies, one reaction without primary antibody was included as a negative control (Supplemental figure 3B, D and F).

### Vitamin A, vitamin E and mineral concentration in liver and serum

Copper, iron, zinc, manganese, molybdenum, cobalt, arsenic, cadmium, lead, thallium, selenium, vitamin E, retinol, palmate and total vitamin A concentrations were determined on 49 of the 86 frozen liver samples by inductively coupled plasma mass spectrometry (ICP-MS) and chromatography at Texas A&M Veterinary Medical Diagnostic Laboratories. The same laboratory determined copper, iron, zinc, manganese, molybdenum, cobalt, arsenic, cadmium, lead, thallium, selenium, vitamin E concentration on 42 samples of frozen serum. Metals and minerals were all assayed on ICP-MS, each element with its own standard curve of at least 5 points with correlation coefficient greater than 0.995. Vitamin A, retinol, palmate and vitamin E were measured by chromatography performed on a Waters system using a C18 Cosmosil PBr column (4.6×250mm). Detailed protocols for ICP-MS and chromatography are provided in the supplementary material.

### Statistical analysis

Equality of means between the two groups of data was assessed using *t* test and PRISM 9 was used for statistic calculations and graphs design.

## Supporting information

Supplemental tables, figures and protocols

## Acknowledgements

The authors would like to thank Lynn Stevenson and Frazer Bell, Diagnostic Services, MVLS, School of Veterinary Medicine, University of Glasgow; Marcus Reddell, Tallowcreek Ranch, Texas; Susan Peters and Brian Cloak, University College Dublin, School of Veterinary Medicine for their technical support. The authors also wish to thank Dr Catherine Barr, Maritza Anguiano, Texas Veterinary Medical Diagnostic Laboratory and Dr Peter O’Brien, University College Dublin, School of Veterinary Medicine for their contributions to the clinical investigation. We also wish to thank Prof. Lorenzo Alibardi, Dipartimento di Scienze Biologiche, Geologiche e Ambientali Università di Bologna, for his support on the development of immunohistochemical protocols.

## Author contributions

P.A.K. and D.S. and A.B. conceived the study. P.A.K and I.M.P designed the study with the support of D.S and A.B. P.A.K., I.M.P and J.D.D carried out the histopathological examination and interpretation of tissue samples. A.B. and D.S. examined the alligators on farm, collected samples, contributed photos and data from the animals included in this study. I.M.P. analyzed the data with the support of D.S and A.B. I.M.P. wrote the manuscript. J.D.D and P.A.K contributed to the writing and revised the manuscript. All authors reviewed the manuscript and consented to its submission and publication.

