## Supplemental tables, figures and protocols for "Hepatic fibrosis induced zinc-deficient dermatosis in American alligators (*Alligator mississippiensis*)"

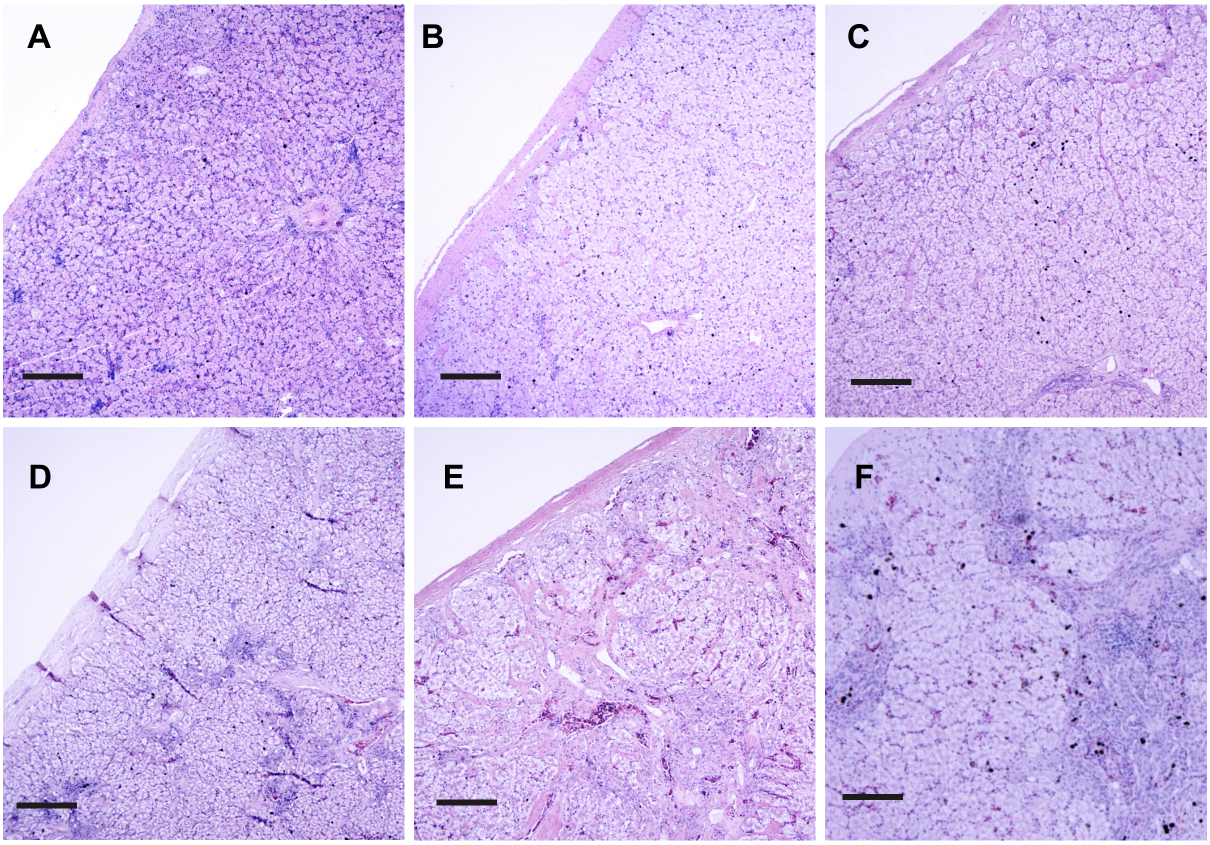


**Supplemental Figure 1. Severity of liver fibrosis observed in alligator livers.** Microphotographs of alligator livers. H&E. Scale bar 1mm. Normal liver (A), liver with mild fibrosis (B), mild to moderate fibrosis (C), moderate fibrosis (D), moderate to severe fibrosis (E) and severe fibrosis (F).

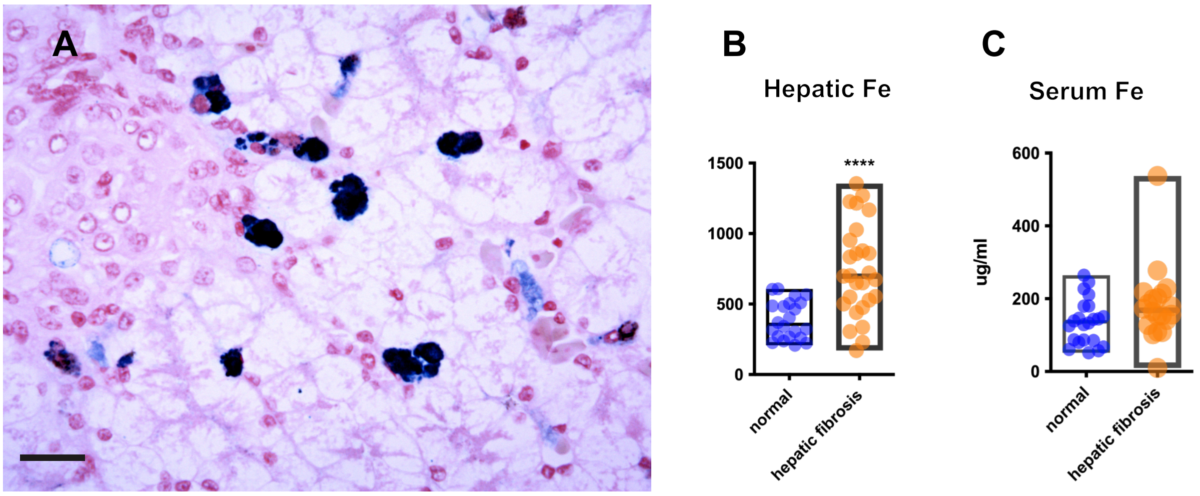


**Supplemental figure 2. Iron loading in association to liver fibrosis, American alligator.** (A) Microphotographs of liver with fibrosis. Perl Prussian Blue. Scale bar 200*μ*m. Iron deposits within macrophages are scattered mainly in the periportal areas. Note the hepatocytes are negative for iron. Box graphs show levels of hepatic (B) and serum (C) iron in alligators with liver fibrosis (n=28) compared to alligators with normal livers (n=22). Central line in the box indicates mean. Difference of means calculated with Student *t* test was significant with $p$ value <0.0001 (****).

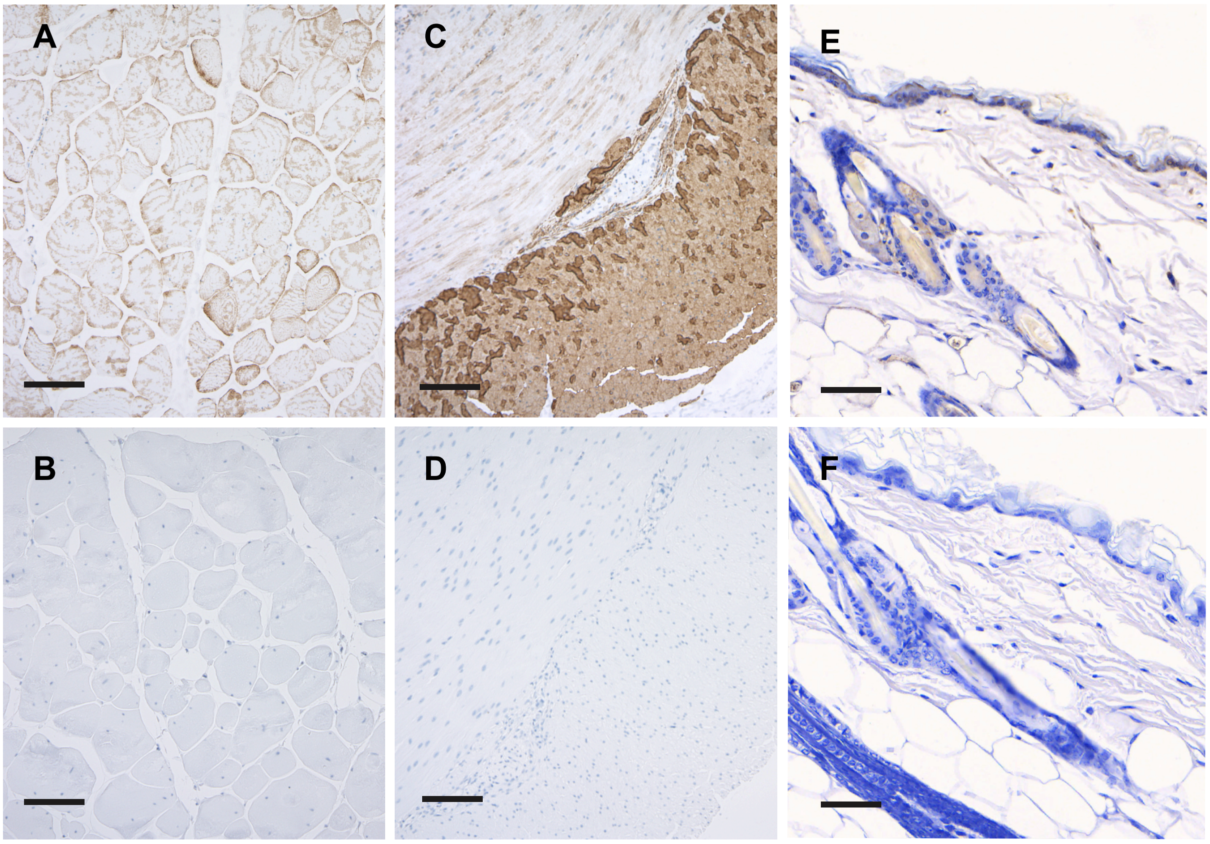


**Supplemental Figure 3. Positive and negative controls for IHC in alligator tissues.** Alligator skeletal muscle labelled with anti-desmin antibody (A) and its negative control (B). Alligator intestine labelled with $\alpha$-SMA-positive antibody (C) and its negative control (D). Canine skin labeled with anti-metallothionein antibody (E). Note the difference in labelling between positive and negative controls (F). Scale bar 200$\mu$m.

**Inductively Coupled Plasma Mass Spectrometry (ICP-MS) protocol**

Metals and minerals were all assayed on ICP-MS, each element with its own standard curve of at least 5 points with correlation coefficient greater than 0.995. All samples were diluted using an internal standard solution whose performance was monitored. NIST Bovine Liver Standard Reference Material and NRCC Dogfish Liver Certified Reference material were assayed alongside each sample batch and their performances monitored for each element. Elemental concentrations were reported on a dry matter basis.

Alligator liver samples whose mass was measured to the nearest 0.0000 g were dried to constant weight in a convection oven at 95°C and reweighed to determine dry weight fraction. For each animal a liver aliquot of ~ 0.5 g was digested using 5.00 mL ultrapure nitric acid, 0.25 mL ultrapure hydrochloric acid and addition of 1 mL 30% hydrogen peroxide. Samples were diluted to 50 mL with tissue diluent containing internal standard and were transferred to the ICP-MS.

**Chromatography protocol**

Vitamin A was measured by chromatography was performed on a Waters system using a C18 Cosmosil PBr column (4.6x250mm) and mobile phases acetonitrile and 0.1 % acetic acid in a gradient with UV monitoring at 350 nm for vitamin A. In a darkened room and BSC, for each animal a liver aliquot of ~1 g (measured to the nearest 0.01 g) was homogenized in de-gassed, deionized water with a final added volume of 5 mL. Betahydrohybutryrate in ethanol was added and samples were subjected continuous rotation at ~30 rpm for 20 minutes. Each tube then received 2 mL hexane and was recapped and rotated for another 20 minutes. After centrifugation at ~2500 rpm to ensure separation of phases, 1 mL of each hexane layer was evaporated to dryness under nitrogen. Resuspension in acetonitrile was followed by filtration at 0.45 um into chromatography vials.

Inductively Coupled Plasma Mass Spectrometry (ICP-MS) and chromatography at Texas A&M Veterinary Medical Diagnostic Laboratories.

**Supplemental Table 1. Samples included in the study**

| sample no | sample collection date | age in months | skin sample | ICP-MS serum sample | ICP-MS liver sample | fibrosis | bile duplication | hepatocyte vacuolation | total histological score |
|---|---|---|---|---|---|---|---|---|---|
| 1 | Jun-19 | 12 | | | | 4 | 5 | 3 | 12 |
| 2 | Jun-19 | 12 | | | | 3 | 3 | 2 | 8 |
| 3 | Jun-19 | 12 | | | | 2 | 2 | 1 | 5 |
| 4 | Jun-19 | 12 | | | | 0 | 2 | 1 | 3 |
| 5 | Jun-19 | 12 | | | | 0 | 2 | 1 | 3 |
| 6 | Jun-19 | 11 | yes,DS | | yes | 3 | 3 | 1 | 7 |
| 7 | Jun-19 | 11 | | | yes | 5 | 5 | 5 | 15 |
| 8 | Apr-19 | 9 | yes,RS | | | 4 | 4 | 3 | 11 |
| 9 | Apr-19 | 9 | yes,RS | | | 2 | 2 | 2 | 6 |
| 10 | Apr-19 | 9 | yes,RS | | | 1 | 2 | 2 | 4 |
| 11 | Jun-19 | 11 | yes,DS | | yes | 4 | 4 | 3 | 11 |
| 12 | Jun-19 | 11 | yes,DS | | yes | 1 | 2 | 3 | 7 |
| 13 | Jun-19 | 11 | | | yes | 1 | 1 | 2 | 4 |
| 14 | Jun-19 | 11 | | | yes | 0 | 1 | 1 | 2 |
| 15 | Jun-19 | 11 | | | yes | 1 | 2 | 1 | 4 |
| 16 | Jun-19 | 11 | | | yes | 0 | 2 | 1 | 3 |
| 17 | Jun-19 | 11 | yes,DS | | yes | 3 | 3 | 1 | 7 |
| 18 | Jun-19 | 11 | | | | 1 | 2 | 1 | 4 |
| 19 | Feb-20 | 19 | | yes | yes | 3 | 4 | 3 | 10 |
| 20 | Feb-20 | 19 | yes,DS | yes | yes | 5 | 5 | 2 | 12 |
| 21 | Feb-20 | 19 | | yes | yes | 2 | 2 | 1 | 5 |
| 22 | Feb-20 | 19 | yes,DS | yes | yes | 1 | 2 | 1 | 3 |
| 23 | Feb-20 | 19 | yes,RS | yes | yes | 1 | 1 | 2 | 4 |
| 24 | Feb-20 | 19 | yes-RS | yes | yes | 5 | 5 | 5 | 15 |
| 25 | Feb-20 | 19 | yes,RS | yes | yes | 3 | 5 | 2 | 10 |
| 26 | Feb-20 | 19 | yes-RS | yes | yes | 3 | 3 | 2 | 8 |

| | | | | | | | | | |
|---|---|---|---|---|---|---|---|---|---|
| 27 | Feb-20 | 19 | yes,DS | yes | yes | 2 | 2 | 1 | 5 |
| 28 | Feb-20 | 19 | yes,DS | yes | yes | 5 | 5 | 3 | 13 |
| 29 | Jan-20 | 18 | | | yes | 0 | 1 | 1 | 2 |
| 30 | Jan-20 | 18 | | | yes | 0 | 1 | 1 | 2 |
| 31 | Jan-20 | 18 | | yes | yes | 0 | 1 | 2 | 2 |
| 32 | Jan-20 | 18 | | yes | yes | 0 | 1 | 2 | 3 |
| 33 | Jan-20 | 18 | | yes | yes | 0 | 1 | 2 | 3 |
| 34 | Jan-20 | 18 | | yes | yes | 0 | 1 | 1 | 2 |
| 35 | Jan-20 | 18 | | yes | yes | 0 | 2 | 1 | 3 |
| 36 | Jan-20 | 18 | | yes | yes | 0 | 1 | 2 | 3 |
| 37 | Jan-20 | 18 | | yes | yes | 0 | 2 | 1 | 3 |
| 38 | Jan-20 | 18 | | yes | yes | 0 | 2 | 1 | 3 |
| 39 | Jan-20 | 18 | | yes | yes | 1 | 2 | 1 | 4 |
| 40 | Jan-20 | 18 | | yes | yes | 2 | 2 | 1 | 5 |
| 41 | Jan-20 | 18 | | yes | yes | 1 | 2 | 2 | 5 |
| 42 | Jan-20 | 18 | | yes | yes | 4 | 5 | 3 | 12 |
| 43 | Jan-20 | 18 | yes,RS | yes | yes | 5 | 5 | 3 | 13 |
| 44 | Jan-20 | 18 | yes-RS | yes | yes | 5 | 5 | 3 | 13 |
| 45 | Jan-20 | 18 | | yes | yes | 5 | 4 | 2 | 11 |
| 46 | Jan-20 | 18 | | yes | yes | 4 | 4 | 2 | 10 |
| 47 | Jan-20 | 18 | | yes | yes | 3 | 3 | 1 | 6 |
| 48 | Jan-20 | 18 | yes,DS | yes | yes | 2 | 2 | 1 | 5 |
| 49 | 13-May-20 | 10 | | | | 0 | 0 | 0 | 0 |
| 50 | 13-May-20 | 10 | | yes | yes | 0 | 0 | 0 | 0 |
| 51 | 13-May-20 | 10 | | yes | yes | 0 | 0 | 0 | 0 |
| 52 | 13-May-20 | 10 | | yes | yes | 0 | 1 | 0 | 1 |
| 53 | 13-May-20 | 10 | | yes | yes | 0 | 0 | 0 | 0 |
| 54 | 13-May-20 | 10 | | yes | yes | 0 | 0 | 0 | 0 |
| 56 | May-20 | 10 | yes,N | | | 0 | 0 | 0 | 0 |
| 57 | May-20 | 10 | yes,N | | | 0 | 1 | 0 | 1 |
| 58 | May-20 | 10 | yes,N | | | 0 | 1 | 0 | 1 |
| 59 | May-20 | 10 | yes,N | | | 0 | 0 | 0 | 0 |

| | | | | | | | | | |
|---|---|---|---|---|---|---|---|---|---|
| 60 | May-20 | 10 | yes,N | | | 0 | 0 | 0 | 0 |
| 61 | May-20 | 10 | | yes | yes | 0 | 0 | 0 | 0 |
| 62 | May-20 | 10 | | yes | yes | 0 | 0 | 0 | 0 |
| 63 | May-20 | 10 | | yes | yes | 0 | 0 | 0 | 0 |
| 64 | May-20 | 10 | | yes | yes | 0 | 0 | 0 | 0 |
| 65 | May-20 | 10 | yes,N | | | 0 | 1 | 0 | 1 |
| 66 | May-20 | 10 | yes,N | | | 0 | 0 | 1 | 1 |
| 67 | May-20 | 10 | yes,N | | | 0 | 0 | 0 | 1 |
| 68 | May-20 | 10 | yes,N | | | 0 | 0 | 1 | 1 |
| 69 | 15-Jun-20 | 11 | | | | 0 | 0 | 0 | 0 |
| 70 | 15-Jun-20 | 11 | | yes | yes | 0 | 0 | 0 | 0 |
| 71 | 15-Jun-20 | 11 | | yes | yes | 0 | 0 | 0 | 0 |
| 72 | 15-Jun-20 | 11 | | yes | yes | 0 | 0 | 0 | 0 |
| 73 | 15-Jun-20 | 11 | | yes | yes | 0 | 0 | 0 | 0 |
| 74 | 15-Jun-20 | 11 | | | | 0 | 0 | 0 | 0 |
| 75 | 15-Jun-20 | 11 | | yes | yes | 0 | 0 | 0 | 0 |
| 76 | 15-Jun-20 | 11 | | yes | yes | 0 | 0 | 0 | 0 |
| 77 | 13-Jun-20 | 11 | | yes | yes | 0 | 0 | 0 | 0 |
| 78 | 13-Jun-20 | 11 | | yes | yes | 0 | 0 | 0 | 0 |
| 79 | 13-Jun-20 | 11 | | yes | yes | 0 | 0 | 0 | 0 |
| 80 | 13-Jun-20 | 11 | | yes | yes | 0 | 0 | 0 | 0 |
| 81 | 13-Jun-20 | 11 | | | | 0 | 0 | 0 | 0 |
| 82 | 13-Jun-20 | 11 | | yes | yes | 0 | 0 | 0 | 0 |
| 83 | 13-Jun-20 | 11 | | | | 0 | 0 | 0 | 0 |
| 84 | 13-Jun-20 | 11 | | yes | yes | 0 | 0 | 0 | 0 |
| 85 | 13-Jun-20 | 11 | | | | 0 | 0 | 0 | 0 |
| 86 | 13-Jun-20 | 11 | | yes | yes | 0 | 0 | 0 | 0 |

DS duoble scale, N normal scale

Fibrosis, biliary duplication and hepayocyte vacuolation:  mild=1, mild to moderate=2, moderate=3, moderate to marked=4, marked=5

**Supplemental table 2. Average body length and belly width of alligator per age group.**

| months of age | body length in cm | belly width in cm |
|---|---|---|
| 9 | 101 | 22 |
| 12 | 110 | 23 |
| 24 | 147 | 30 |
| 36 | 184 | 37 |
